# Label-free tumor cells classification using deep learning and high-content imaging

**DOI:** 10.1101/2023.02.03.526929

**Authors:** Chawan Piansaddhayanon, Chonnuttida Koracharkornradt, Napat Laosaengpha, Qingyi Tao, Praewphan Ingrungruanglert, Nipan Israsena, Ekapol Chuangsuwanich, Sira Sriswasdi

**Affiliations:** Department of Computer Engineering, Faculty of Engineering, Chulalongkorn University; Center of Excellence in Computational Molecular Biology, Faculty of Medicine, Chulalongkorn University; Chula Intelligent and Complex Systems, Faculty of Science, Chulalongkorn University; NVIDIA AI Technology Center, Singapore; Center of Excellence for Stem Cell and Cell Therapy, Faculty of Medicine, Chulalongkorn University; Department of Pharmacology, Faculty of Medicine, Chulalongkorn University; Center for Artificial Intelligence in Medicine, Research Affairs, Faculty of Medicine, Chulalongkorn University

## Abstract

Many studies have shown that cellular morphology can be used to distinguish spiked-in tumor cells in blood sample background. However, most validation experiments included only homogeneous cell lines and inadequately captured the broad morphological heterogeneity of cancer cells. Furthermore, normal, non-blood cells could be erroneously classified as cancer because their morphology differ from blood cells. Here, we constructed a database of microscopic images of organoid-derived cancer and normal cell with diverse morphology and developed a proof-of-concept deep learning model that can distinguish cancer cells from normal cells within an unlabeled microscopy image. In total, more than 75,000 organoid-drived cells from 3 cholangiocarcinoma patients were collected. The model achieved an area under the receiver operating characteristics curve (AUROC) of 0.78 and can generalize to cell images from an unseen patient. These resources serve as a foundation for an automated, robust platform for circulating tumor cell detection.

## Background & Summary

Circulating tumor cell (CTC), or cell from primary tumor that were shed into the patient’s bloodstream, holds important clinical values as a source of early, non-invasive biomarker of metastasis and cancer prognosis and many cancer types (1, 2). Existing technologies for isolating and detecting CTC mainly rely on the fact that most normal blood cells can be captured by antibody targeting certain cell surface markers, such as CD45, while tumor cells can be captured by antibody targeting different markers (3). Although multiple antibodies have been developed for characterizing various CTC types, such as epithelial and mesenchymal CTC (4), enrichment-based approaches still cannot account for the full heterogenicity of CTC. In fact, a study of lung cancer patients has shown that only 40-60% of CTC in blood samples were detected by enrichment-based approaches (5). Nowadays, high-throughput sequencing technologies have also been applied to characterize the genome and transcriptome of individual CTC (6) as a non-invasive mean to probe the molecular signature of primary tumors and to develop prognostic cancer biomarkers.

Another possibility for unbiased characterization of individual CTC is through high-content microscopy imaging of patient blood samples, whereby cancer cells can be differentiated from normal cells as well as classified into types based on their distinctive morphological properties (7, 8). These techniques are enabled by recent advances in deep learning which let us train artificial neural network models to accurately identify cell types (9, 10) and pinpointing the locations of subcellular compartments (11, 12) from bright-field microscopy images without any labeling of the cells. However, imaging-based CTC detections were mostly developed and/or validated only on spiked-in cells from a few cell lines that do not capture the broad heterogeneity and morphological properties of actual CTC (13). For example, Wang *et al*.(14) trained a deep learning model using 436 cultured cells and 1,309 white blood cells and validated their model on 32 CTCs from two patients. A recent work has also shown that CTCs derived from different tumor sites exhibit clearly distinct morphological characteristics (15). This suggests the possibility of simultaneously detecting and predicting the tissue-of-origin for each CTC.

Hence, the first step toward developing a generalized imaging-based CTC detection platform is to establish a large-scale microscopy imaging database of cancer and normal cells that capture the heterogeneity of both cancer types and tissue types. Patient-derived organoids, or 3D cultures, have been shown as realistic sources of diverse cell types and morphology that faithfully represent the genotype and phenotype of cancer subtypes (16, 17). The combination of paired cancer and normal cells derived from the same tissue of the same patient would serve as a good benchmark for an imaging-based CTC detection technique by testing whether the technique can distinguish between cancer and normal cells (as supposed to distinguishing between blood and non-blood cells). By expanding the database of cell images to cover multiple tissues, cancer types, and patients, and by linking cell images to prognosis and treatment response information, future imaging-based CTC platforms have the potential to not only detect CTC, but also predict the tissue-of-origin and aid clinical decision making.

In this research, a large database of microscopic images of more than 75,000 individual organoid-derived cancer and normal cells from 3 cholangiocarcinoma patients were constructed, and a proof-of-concept deep neural network model was developed to (i) evaluate the possibility of distinguishing cancer and normal cells based on only unlabeled bright-field microscopic images and (ii) explore the morphological diversity of cancer and normal cells across cancer types and individual patients. The full dataset and code used for development are available at https://figshare.com/articles/dataset/Fluorescence_imaging_of_CCA_organoid-derived_cells/19960232 and https://github.com/cmb-chula/CancerCellVision-CCA, respectively.

## Methods

### Cholangiocyte organoid culture

Liver biopsies were cut into small pieces and washed 3 times with Advanced DMEM/F12 supplemented with 1x Gluta-max, 10 mM HEPES, and 1x antibiotics (AdDF+++, Gibco, Thermo Scientific). Liver tissues were digested using 100 *μ*g/ml dispase I and 300 U/ml collagenase XI in Cholangiocyte culture media with Advanced DMEM/F12 containing 10% R-Spondin condition media, 10% Wnt3a condition media, 1 mM N-Acetylcysteine, 10 mM Nicotinamide, 1× B27 supplement, 1× N2 supplement, 100 ng/ml Noggin, 10 nM Gastrin-I, 50 ng/ml EGF, 5 uM A83-01, 100 ng/ml FGF10 (Peprotech), 25 ng/ml HGF (R&D Systems), and 10 *μ*M FSK (Tocris). The cultures were incubated at 37°C for 1 hour. The digestion reaction was stopped with 10 ml AdDF+++ and the resulting suspension was filtered through a 70 *μ*M cell strainer. Cells in suspension were collected via centrifugation and washed 5 times with AdDF+++. Cell pellets were resuspended in 70% Matrigel (Corning) and dropped on pre-warmed 24-well culture plates. After the Matrigel solidified, 500 *μ*l of organoid culture media was added. Cells were cultured at 37°C with 5% CO2. The media were changed every 3 days and the cell passage was performed every 1-2 weeks by mechanically dissociating the cells with P1000 pipette tip.

### Fluorescence labeling and high-content imaging

Each organoid was dissociated into single cells using TrypLETM Express Enzyme (Gibco, Thermo Scientific). Around 106 cancer and normal cells were obtained from each sample. Cells from cancer organoids were stained with a deep red fluorescence (Cytopainter ab176736) while cells from normal organoids were stained with green fluorescence (Cytopainter ab176735). Nuclei were stained with Hoechst. Cancer and normal cells were mixed at 1:1 ratio, dropped on 96-well plates, and subjected to bright-field and fluorescence imaging on an Opera Phenix instrument (Perkin Elmer). In total, 1207 paired bright-field and fluorescence images were acquired for cancer and normal cholangiocytes. Each image consists of 1080×1080 pixels and contains 20-30 individual cells on average.

### Image processing and preparation

Prior to the annotation step, brightfield and fluorescence images were prepossessed to make the individual cells more visually distinguishable to the human eyes. Data preprocessing steps described in Christiansen *et al*. (11) were performed with some modification. First, a median filter of size 5 × 5 was repeatedly applied to the fluorescence images until convergence to reduce the salt-and-pepper noise. After that, images were bilinearly downsampled by a factor of two to reduce shot noise. Finally, pixel intensities were normalized per image to the same mean and standard deviation. Frame stitching did not need to be performed due to the difference in data acquisition technique. Flat field correction and dust artifact removal were also not applied because these operations did not significantly affect the quality of images here. After preprocessing, the three fluorescence activations (red for cancer cells, green for normal cells, and Hoechst blue for nuclei) of each image were merged into a single three-channel image. Examples of prepossessed and annotated images are shown in Fig 1.

**Figure 1:**
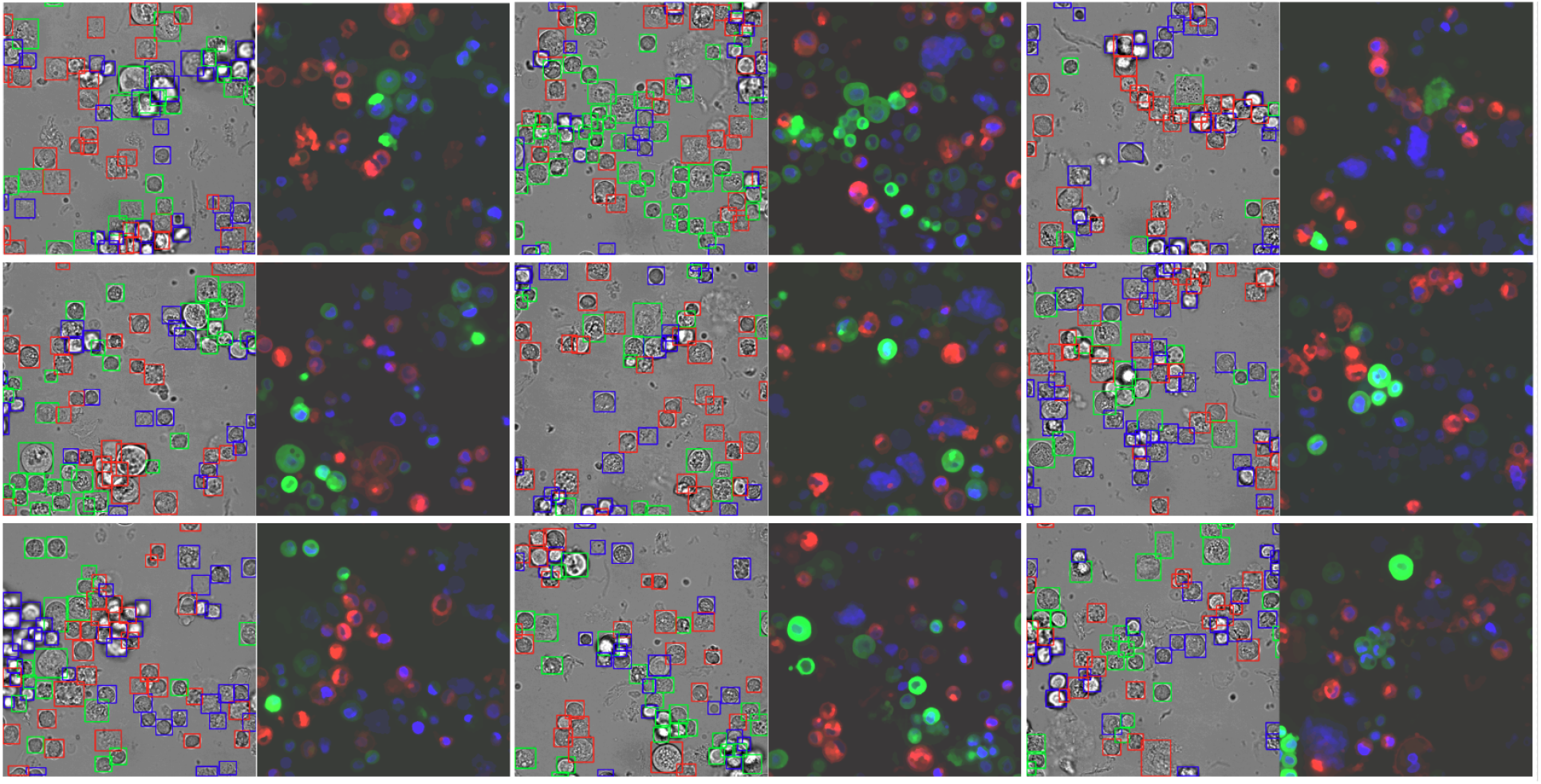
Examples of preprocessed and annotated brightfield and fluorescence image for human annotator. Box colors indicate the object classes (red for cancer cells, green for normal cells, and blue for unknown cells that exhibited neither signals).

### Cell annotation

There were three human annotators. One annotator is an expert in microscopy with more than three years of experience. The other annotators are graduate biology students. An inter-annotator agreement was evaluated at the beginning by asking all three annotators to analyze the same set of 6 images (about 150 individual cells). Labelme (18) was used to annotate the location and classification of each cell. Brightfield image and the corresponding fluorescence image were simultaneously shown to the annotators. Cells were classified as either cancer, if there was a clear red fluorescence signal, normal, if there was a clear green fluorescence signal, or unknown, if only the Hoechst signal was visible (Fig 2). bounding boxes and classification for the remaining images and the results were provided to the annotators for further refinements. Annotators can add new bounding boxes, remove erroneous bounding boxes, or change the classification of each cell. At the end of the second phase, 1087 out of 1207 images were analyzed by at least one annotator. These data were used to train the proof-of-concept model.

**Figure 2:**
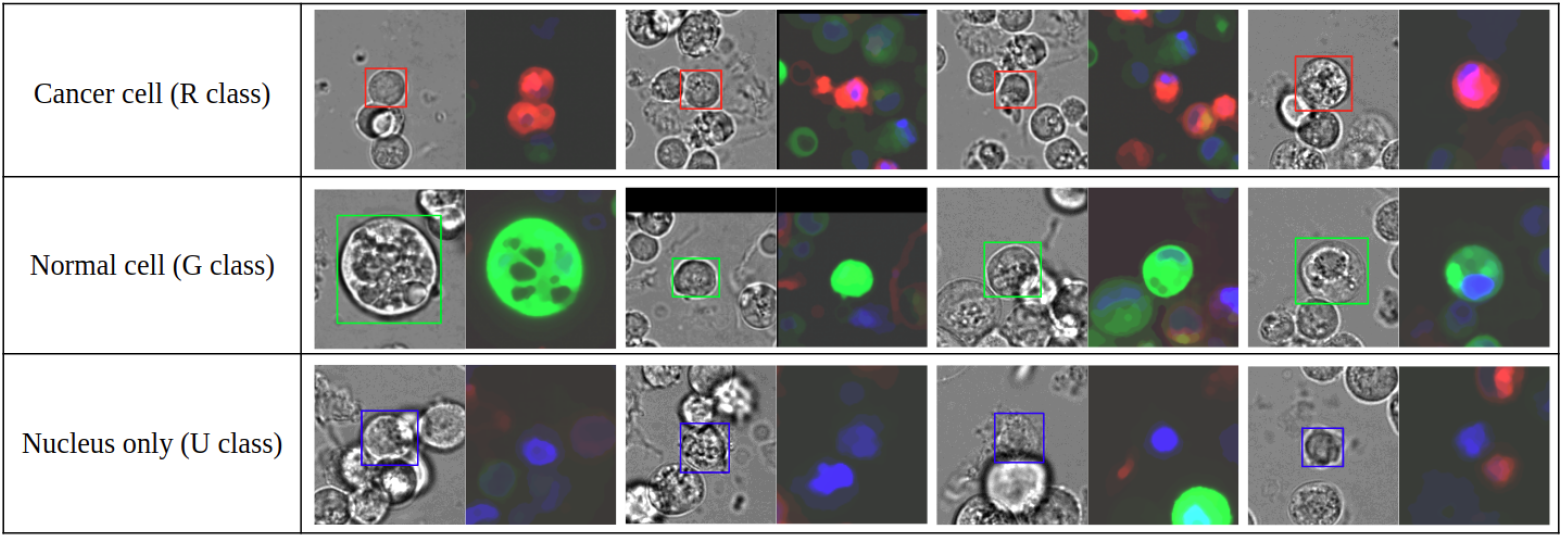
Examples of annotated cells from each class.

The annotation process were divided into three phases (Fig 3 and Fig 4). In the first phase, a subset of 30 images were fully annotated by the most experienced annotator and then used to train an initial object detection model, with both brightfield and fluorescence images as inputs. In the second phase, the initial model was used to generate

**Figure 3:**
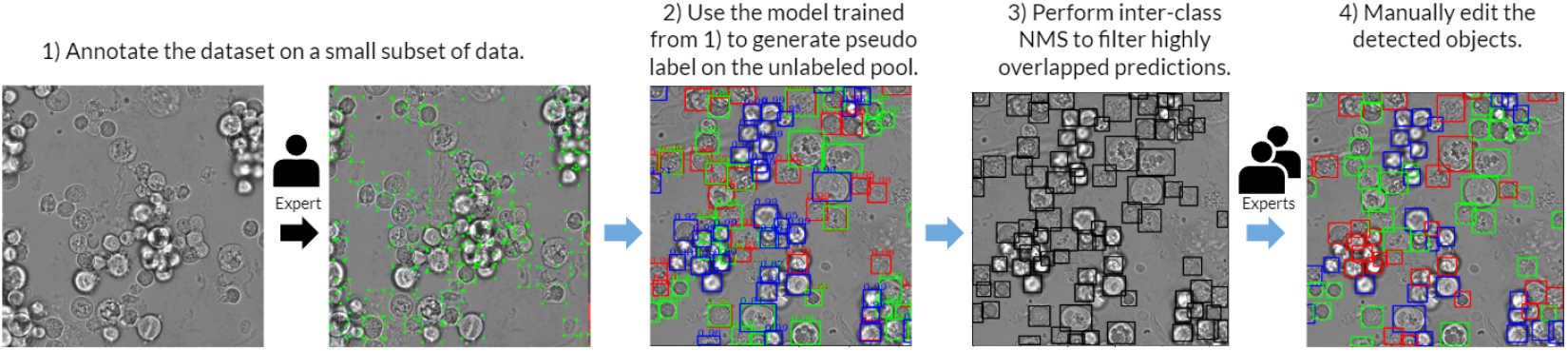
The annotation process for the training and validation sets. First, a small subset (30 images) was fully manually annotated. Then, the initial cell detection and classification model was trained to generate pseudolabels for all unannotated images. The pseudo-generated bounding boxes were then filtered using Non-Maximum Suppression (NMS) to remove highly overlapping boxes. These pseudolabel annotation were then refined by the experts to obtain the final annotation used for training and validation. Note that every step in this annotation process used fluorescence images as guidance.

**Figure 4:**
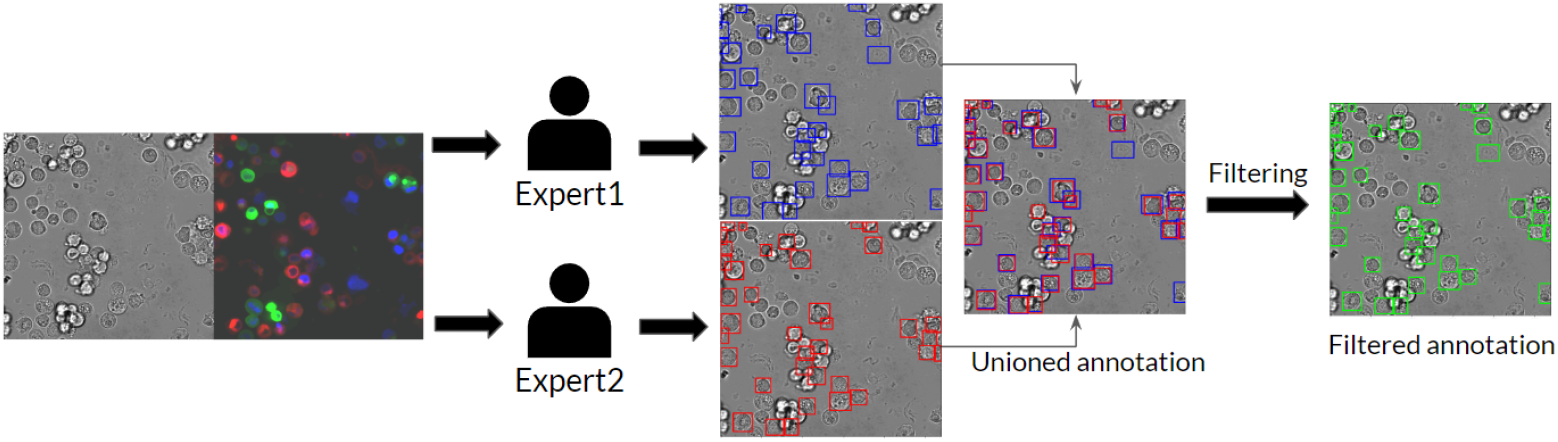
The annotation process for the test set. Two annotators were separately tasked to annotate all cancer cells inside each brightfield image with paired fluorescence image as guidance. Results from the two annotators were combined and used as the final annotation.

In the third phase, to construct the test set, 120 images were sampled from three patients (40 images each) and manually annotated by human annotators. Bounding boxes and classification labels from the initial object detection model were intentionally withheld to minimize biases. Furthermore, to maintain high annotation quality, each image was analyzed by at least two annotators and only cancer cells were annotated. The bounding boxes defined by the two annotators were merged using non-maximum suppression (NMS). When there is disagreement, bounding boxes produced by the annotator with more experience were used.

### Data Records

The dataset consists of 1207 paired brightfield and fluorescence microscopy images with a resolution of 1080 *×* 1080 in the TIFF format with cell-level bounding box and classification annotations in the VOC format. There are 84,503 cell-level bounding box annotations consisting of a bounding box (xmin, ymin, w, h), and object class. The three object classes are R, G, and U, which refer to tumor cell (red fluorescence), normal cell (green fluorescence), and unknown cell, respectively. The dataset is separated into training, validation, and test splits, where the test split contains only cancer cell annotation, while the rest have all three classes. The number of objects from each class in each data split is shown in Table 1. The dataset is available on FigShare at https://figshare.com/articles/dataset/Fluorescence_imaging_of_CCA_organoid-derived_cells/19960232.

**Table 1:** The number of images and cells in each dataset split.

| Data split | No. of image | R | G | U |
| --- | --- | --- | --- | --- |
| Train | 967 | 22800 | 24862 | 24575 |
| Validation | 120 | 2823 | 2972 | 3135 |
| Test | 120 | 3336 | - | - |

### Detailed description

Figure 5 summarizes the indexing structure of our dataset. Original raw image files are stored in the directory `raw_images_for_model`. This directory consists of two sub-directories: `raw_images_for_model/brightfield` contains brightfield images and `raw_images_for_model/fluorescence` contains fluorescence images. Files are named with the `r{patient_id}c04f{file_id}p01.tiff` format, where `patient_id` and `file_id` refers to the IDs of the patients (06, 07 or 08) and image, respectively. Each brightfield image and the corresponding fluorescence image share the same file name. Each fluorescence image is a three-channel image file where channels correspond to red fluorescence signal (cancer cells), green fluorescence signal (normal cells), and Hoechst signal (nuclei), respectively. These raw images can be readily used as input for the detection stage without further post-processing.

**Figure 5:**
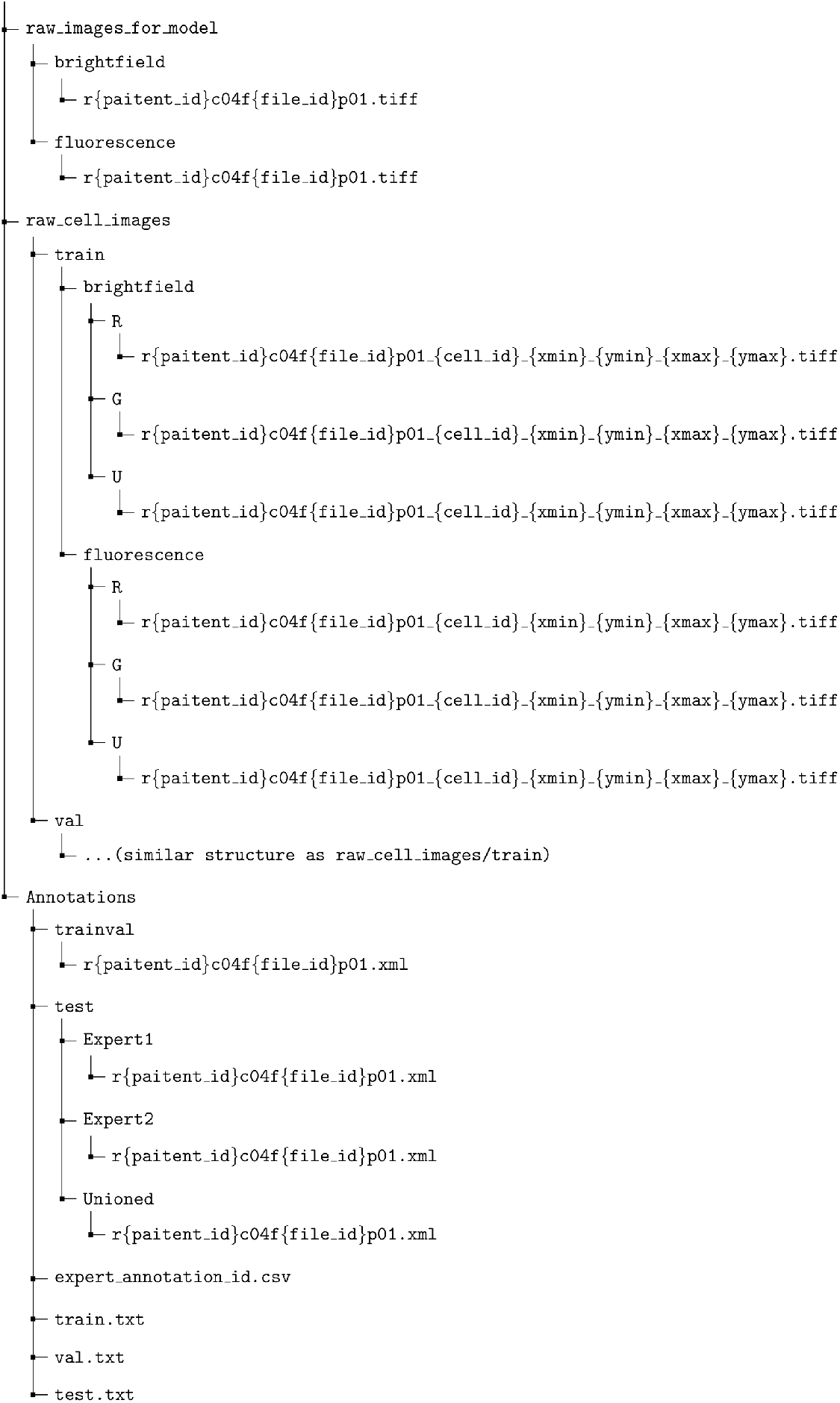
The index of our proposed dataset.

Annotations are provided in the directory `Annotations`. Annotations for the training-validation split and the test split are provided separately in subdirectories `trainval` and `test`, respectively. Each annotation file is named with the same `r{paitent_id}c04f{file_id}p01.xml` format as the raw image files provided in `raw_images_for_model`. The test subdirectory contains three subdirectories: `Expert1, Expert2`, and `Unioned`, which contain the annotations from the first expert, second expert, and the combined version, respectively.

Images of individual extracted cells, which are ready to use for cell classification, are provided in the directory `raw_-cell_images`. There are three subdirectories `R`, `G` and `U`, each containing images of cells from each class. The file name of each cell follows the `r{paitent_id}c04f{file_id}p01_{cell_id}_{xmin}_{ymin}_{xmax}_-{ymax}.tiff` format, where `cell_id` refers to the ID of each cell, and `(xmin, ymin, xmax, ymax)` indicates the position of the cell in the raw image `r{paitent_id}c04f{file_id}p01.tiff`.

Each line in `train.txt, val.txt`, and `test.txt` indicates the split of each data point. The file `expert_annotation_id.csv` contains the ID of the annotator who analyzed each image.

### Technical Validation

Technical validations of our dataset were conducted by training a deep learning model to recognize cancer cells in given brightfield (unlabeled) microscopy image. Evaluations were performed at two levels: cell level and image level. The cell-level evaluation measures the model’s ability to distinguish between cancer (class R) and other cell types (classes G and U) from given cropped cells from the brightfield image as an input. On the other hand, the image-level evaluation measures the model’s ability to do so on the whole brightfield image. This setup introduces additional challenges since the model also has to differentiate cancer cells from background objects and imaging artifacts.

The experiments were conducted under three input settings: Brightfield, Brightfield + Hoechst, and Brightfield + Fluorescence. The Brightfield setting is a standard setup where the model receives only the brightfield images as an input, while under the Brightfield + Hoechst or Brightfield + Fluorescence settings, Hoechst fluorescence signals or all fluorescence signals were also provided as input, respectively. The Brightfield + Hoechst setting reflects the situation where nuclei staining data are available. The Brightfield + Fluorescence setting was included to evaluate the upper bound of cancer cell recognition performance (as fluorescence signals that contain the ground truth are provided).

Here, a two-stage detection pipeline consisting of a detector and a classifier was developed. The detector is responsible for proposing bounding boxes of objects of interest, while the classifier refines the confidence score of each proposed bounding box. During the cell-level evaluation, the ground truth bounding box of each object was directly provided to the classifier. An overview of the pipeline is shown in Figure 6.

**Figure 6:**
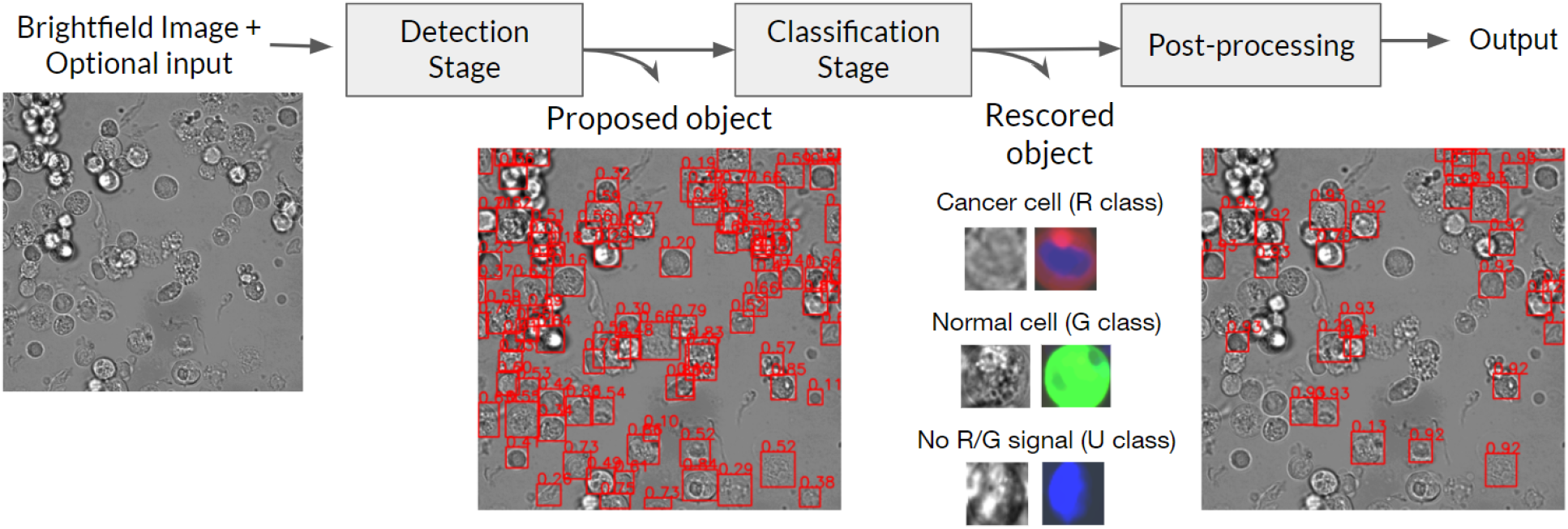
The main pipeline for cancer cell detection consists of two stages, detection and classification, each being a deep artificial neural network. The detector proposes possible cancer cells which are then re-examined by the classifier to refine the confidence scores. Finally, Non-Maximum Suppression (NMS) is performed to remove highly overlapping bounding boxes.

### Cell detection from brightfield image

A deep object detection artificial neural network based on Faster R-CNN (19) with ResNet-50 (20) as a network backbone was developed to propose the locations of all cancer cells in a given brightfield image. The model receives an image as an input and return a set of bounding boxes, {(*x*_1_, *y*_1_, *w*_1_, *h*_1_, *S*_1_), …, (*x_n_*, *y_n_*, *w_n_*, *h_n_*, *S_n_*)}, where each element of a tuple indicates the center of the predicted cell, the dimension of the predicted cell, and the confidence score for the cancer class, respectively. In our benchmarks, the model was trained to detect only cancer cells, as we found that training the model to simultaneously recognize cells from all three classes dampened the performance.

The original 1080 *×* 1080 pixels resolution of the brightfield image was used for training. The network backbone was initialized using ImageNet pre-trained weights (21). Minor modifications were made to adjust the number of output classes and the first convolutional layer. The number of input image channels were adjusted to 4 and 6 accordingly when fluorescence signals are provided as input (the Brightfield + Hoechst and Brightfield + Fluorescence settings). The training framework was based on MMDetection (22). Specifically, the model was trained using a batch size of and stochastic gradient descent (SGD) as an optimizer. The learning rate was set at 10^−3^ for 32 epochs and then divided by a factor of 10 after 16 and 24 epochs have passed. Only random flip augmentation were performed during training.

### Refinement of cell detection results

Downstream from the object detection network is a classifier, which is a deep convolutional neural network (CNN) that outputs the confidence score for each object predicted by the detector. ConvNext-B (23) was used as the network backbone with a fixed input resolution of 128 *×* 128 pixels. The network backbone was initialized using ImageNet (21) pre-trained weights. The model was trained using a batch size of 64 and Adam as an optimizer. The learning rate was set at 5 *×* 10^−4^ for 12,000 iterations and then divided by a factor of 10 after 6,000 and 10,000 iterations have passed. Random geometric augmentation, gaussian blur, and random brightness augmentation were performed during training. During the image-level evaluation, the confidence score *S* is the weighted average between the scores produced by the detector, *S_det_*, and the classifier, *S_cls_*, with the weight *ω*, (*S* = (1 *− ω*)*S_det_* + *ωS_cls_*). *ω* was set to 0 during the cell-level evaluation to disregard the contribution from the detector.

### Cell-level performance evaluation

Cell-level evaluation was performed on three random training-validation splits of the dataset to calculate the mean and standard deviation of each performance metric. The cancer class confidence thresholds that yielded the highest F1 scores were selected for calculating the precision and recall values. The areas under the receiver operating characteristics curve (AUROCs) were also reported.

Table 2 summarized the cell-level performance of our model. Unsurprisingly, when both brightfield and fluorescence images were used as input (the Brightfield + Fluorescence setting), the model could accurately recognize cancer cells with an F1 score of 94.5. While this setting is unrealistic, it confirmed the quality and consistency in the annotations. Figure 8c shows that most of the confusions involved unknown cells, which are either cancer or normal cells that exhibit nuclear staining fluorescence but no cytoplasmic staining fluorescence. There was only around 1% confusion between normal and cancer cells. UMAP visualization (26) of the latent embedding vectors, extracted from the feature map of the last layer before the last global pooling in the neural network, for the individual cells (Figure 7c) shows that unknown cells not only reside between the normal cells and cancer cells but also are visually separable from the other classes.

**Table 2:** Cell-level cancer classification performance of our method on the validation split of our dataset+.

| Setting | F1 | precision | recall | AUROC |
| --- | --- | --- | --- | --- |
| Brightfield | $60.5 \pm 0.4$ | $50.5 \pm 0.6$ | $75.5 \pm 1.7$ | $77.5 \pm 0.2$ |
| Brightfield + Hoechst | $66.0 \pm 0.2$ | $58.7 \pm 0.9$ | $75.5 \pm 1.3$ | $83.0 \pm 0.2$ |
| Brightfield + Fluorescence | $94.5 \pm 0.1$ | $93.5 \pm 0.3$ | $95.5 \pm 0.1$ | $99.3 \pm 0.1$ |

**Figure 7:**
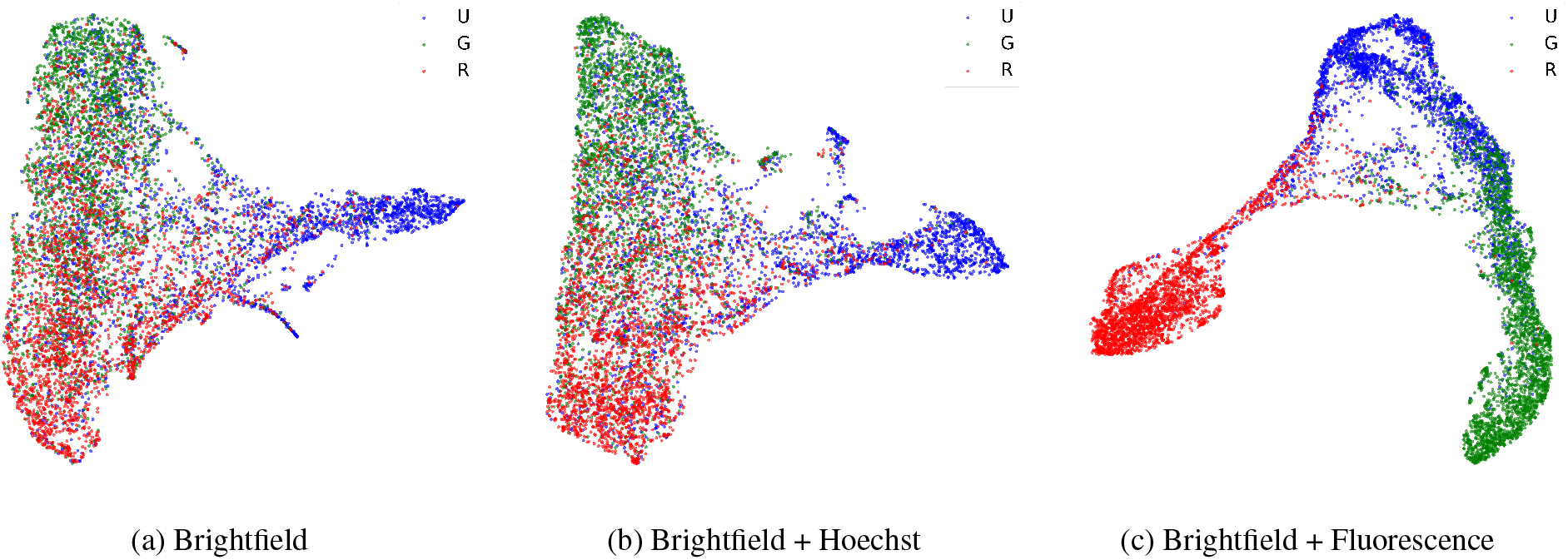
2D embeddings of cells from different classes in the dataset. The embeddings were calculated using UMAP from the feature map at the last layer before the last global average pooling in the network. (a) Embeddings from the model trained using only brightfield images. (b) Embeddings from the model trained with brightfield images and Hoeschst signal. (c) Embeddings from the model trained with brightfield images and all fluorescence signals.

**Figure 8:**
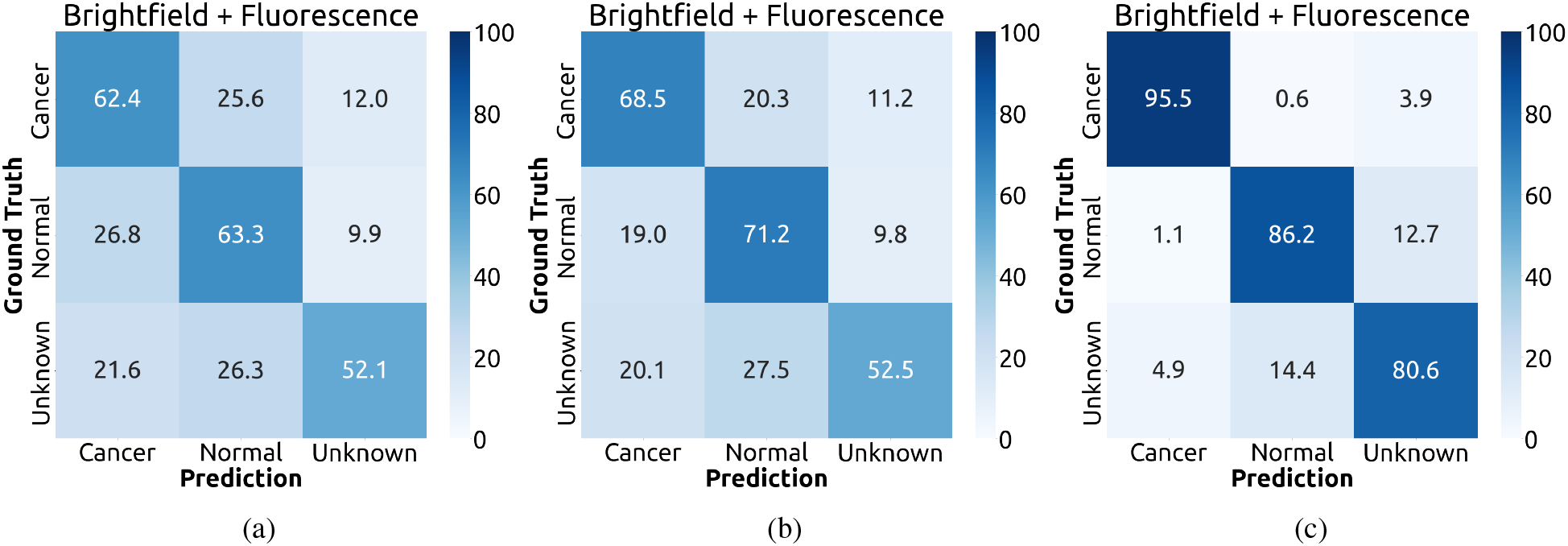
Normalized confusion matrix of the cell-level evaluation on the validation split.

With only brightfield images as input (the Brightfield setting), the cancer cell classification performance dropped to 60.5% F1 with more than 20% confusion between normal and cancer cells (Figure 8a). Visualization of cell embeddings revealed that the boundary between the three cell classes are much more ambiguous (Figure 7a), especially between normal and cancer cells. When the Hoechst fluorescence channel which indicate the nuclei was included as an input, the classification performance improved noticeably to 66.0% F1 (Figure 8b) and the boundary between the three cell classes became clearer (Figure 7b). This indicates that the model can take advantage of the differences in nuclear morphology between normal and cancer cell (27).

To investigate the impact of neural network architecture choice on cancer cell classification performance, an ablation analysis was conducted by replacing the backbone network from ConvNext-B (23) with EfficientNet-B4 (24), DenseNet-121 (25), or ResNet-50 (20). Table 3 indicates that architecture choices can change the performances by up to 2.2% F1 score and 2.1% AUROC, with ConvNext-B achieving the highest performances and ResNet-50 achieving the lowest performances.

**Table 3:** The effect of classifier backbone architecture choices on cell-level performances.

| Network Backbone | Brightfield |  | Brightfield + Hoechst |  | Brightfield + Fluorescence |  |
| --- | --- | --- | --- | --- | --- | --- |
|  | F1 | AUROC | F1 | AUROC | F1 | AUROC |
| ConvNext-B (23) | 60.5 ± 0.4 | 77.5 ± 0.2 | 66.0 ± 0.2 | 83.0 ± 0.2 | 94.5 ± 0.1 | 99.3 ± 0.1 |
| EfficientNet-B4 (24) | 60.0 ± 0.1 | 77.3 ± 0.2 | 65.2 ± 0.3 | 82.2 ± 0.1 | 94.4 ± 0.2 | 99.2 ± 0.1 |
| DenseNet-121 (25) | 59.5 ± 0.2 | 76.5 ± 0.2 | 64.8 ± 0.2 | 81.8 ± 0.1 | 94.2 ± 0.1 | 99.2 ± 0.1 |
| ResNet-50 (20) | 59.1 ± 0.2 | 75.8 ± 0.1 | 63.8 ± 0.2 | 80.9 ± 0.1 | 94.6 ± 0.1 | 99.3 ± 0.1 |

### Image-level performance evaluation

For the image-level evaluation, the ability of the model to locate cancer cells in a large brightfield image is also measured. Each bounding box predicted by the model is considered a match to a cancer cell if it overlaps with the annotated bounding box with an intersection-over-union (IoU) ratio of at least 0.5. Furthermore, because only the cancer cell class is considered here, the average precision at the IoU threshold of 0.5 (AP50) was measured instead of AUROC. F1 scores were also reported for comparison to the cell-level evaluation. Table 4 shows significant performance improvement in both Brightfield and Brightfield + Hoechst settings when the two-stage architecture (full pipeline) was used over the deep object detector (detection stage). This was because the detector can produce high-confidence false positives when many objects overlap with each other, such as in areas with high density of cells. The downstream classification stage can effectively resolve these errors as it observe each proposed object separately. For the Brightfield + Fluorescence setting, the performance did not change much because some of the bounding boxes generated by the detection stage were oversized and did not sufficiently overlap with the ground truth annotation, even though the predicted classes were correct (Fig. 9). It should be noted that a small performance gain can still be achieved by properly weighing the prediction confidences between the detector and the classifier (*ω* = 0.7).

**Table 4:** Image-level cancer cell detection performance of our method on the test split.

| Network Backbone | Detection stage | | Full pipeline ($\omega = 0$) | | Full pipeline ($\omega = 0.7$) | |
| --- | --- | --- | --- | --- | --- | --- |
|  | AP50 | F1 | AP50 | F1 | AP50 | F1 |
| Brightfield (23) | 43.1 ± 0.6 | 50.3 ± 0.3 | 50.6 ± 0.2 | 54.1 ± 0.2 | 52.2 ± 0.2 | 55.4 ± 0.3 |
| Brightfield + Hoechst | 45.5 ± 0.7 | 52.2 ± 0.3 | 56.5 ± 0.5 | 57.8 ± 0.4 | 57.7 ± 0.3 | 59.3 ± 0.3 |
| Brightfield + Fluorescence | 86.6 ± 0.3 | 87.3 ± 0.1 | 86.7 ± 0.3 | 86.2 ± 0.2 | 89.2 ± 0.3 | 87.8 ± 0.1 |

**Figure 9:**
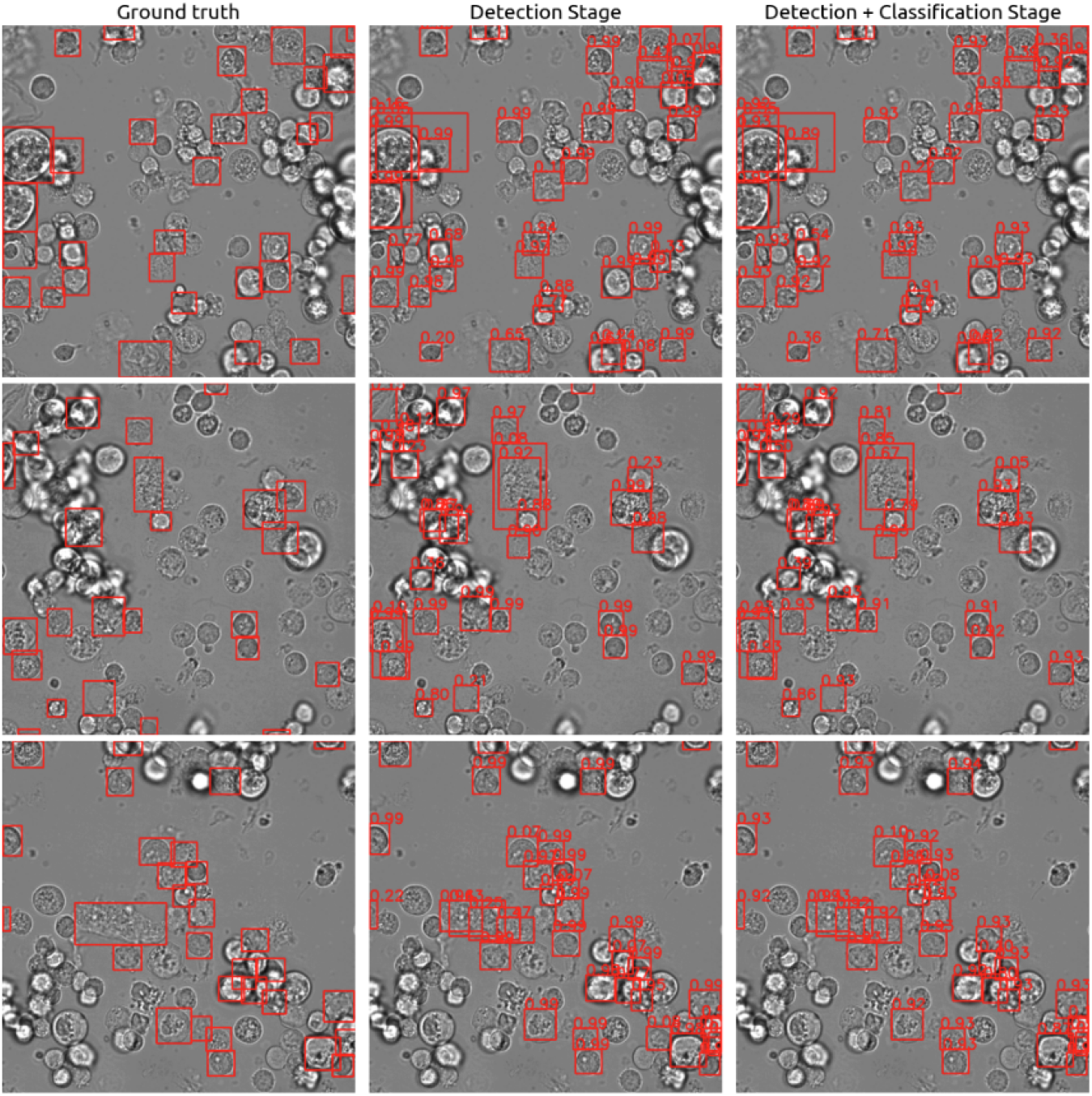
Example of image-level predictions (red boxes) and their confidence on the test set under the Brightfield + Fluorescence setting. Despite having the fluorescence signal as guidance, the model still outputted oversized bounding boxes and could not distinguish individual cells in areas with high cell density.

One interesting result is how information from unknown cells (those with unclear cytoplasmic fluorescence signals) could be used to improve cancer cell detection performance. As shown in Table 5, dropping all unknown cells from the training data resulted in a suboptimal F1 of 56.3%. Thus, we performed semi-supervised learning by predicting pseudolabels for unknown cells and adding them to the training set. However, the performance dropped regardless of whether all pseudolabels were included or even when only high-confidence pseudolabels were considered. Curiously, the best improvement with 3.0% additional F1 was achieved by labeling all unknown cells as non-cancer. This is unexpected because there are many unknown cells whose latent embeddings, which reflect the cells’ morphological characteristics, were similar to cancer cells’ (Figure 7). These unknown cells are expected to be poorly stained cancer cells. A possible explanation is that because the majority of unknown cells are morphologically distinct from both cancer and normal cells (Figure 7), they might include non-cell objects such as dead cells and other debris. Hence, by treating all unknown cells as non-cancer, the model might better delineate the morphological boundary of cancer cells.

**Table 5:**
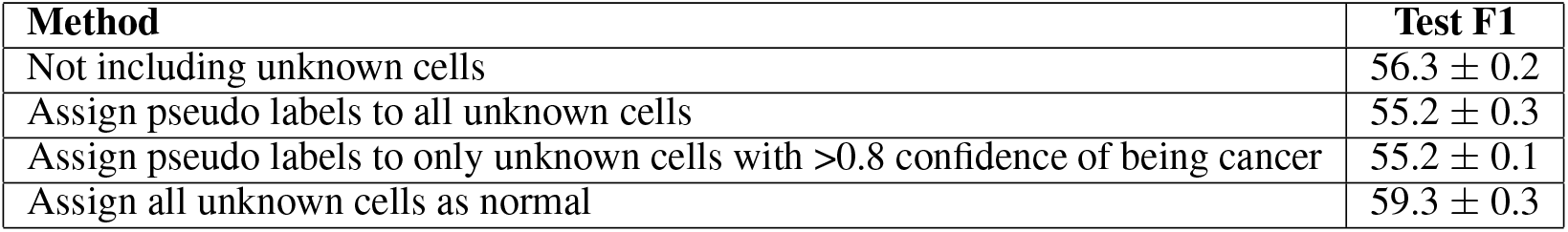
Impact of various strategies for adding unknown cells to the training set on cancer cell detection performance. The experiments were conducted under the Brightfield + Hoechst setting with confidence weighting (*ω* = 0.7) during inference. The confidence of each pseudolabel was obtained from an average of three inference runs from the models of different seeds.

### Evaluation of patient-to-patient variation

The extent of patient-to-patient variation in cell morphology was evaluated by training the model using data from one or two patient(s) and measuring the performance on data from the unseen patient(s). Overall, the model can generalize to cell images from unseen patients with less than 2% drop in F1 (Table 6). Although the performances were lowest when the models were trained or tested on data from the third patient, this is likely because only 91 images from this patient have been annotated by human experts. In contrast, around 500 images were annotated each for the other two patients.

**Table 6:** Model performances (F1) when trained and tested on cell images from different cholangiocarcinoma patients. The experiments were conducted under the Brightfield + Hoechst setting with confidence weighing (*ω* = 0.7) during inference.

| Training data | Number of annotated images | 1<sup>st</sup> patient | 2<sup>nd</sup> patient | 3<sup>rd</sup> patient |
| --- | --- | --- | --- | --- |
| All 3 patients | 967 | 60.1 ± 0.2 | 59.2 ± 0.2 | 59.4 ± 0.5 |
| 1<sup>st</sup> patient | 493 | 58.3 ± 0.2 | 58.3 ± 0.5 | 57.9 ± 0.5 |
| 2<sup>nd</sup> patient | 503 | 57.7 ± 0.2 | 56.9 ± 0.4 | 58.1 ± 0.4 |
| 3<sup>rd</sup> patient | 91 | 39.2 ± 0.4 | 39.7 ± 0.4 | 39.2 ± 0.7 |

### Impact of dataset size on cancer cell classification

Although our dataset already contains 25000-30000 of cells from each class, the broad heterogeneity of cell morphology may not yet be fully captured. To evaluate the impact of additional training data on cancer cell classification, the training set were artificially down-sampled to 5%, 10%, 20%, and 50% of the original size to monitor the gain in performance as the training set size grows. Figure 11 shows that the performance readily saturate with just 5% of the training data if fluorescence signals were provided as input. On the other hand, under realistic settings where brightfield images are the main source of information, cancer cell classification performance increased steadily and linearly as the size of the data grew exponentially. This strongly suggested that the model will benefit from even more training cell images.

**Figure 10:**
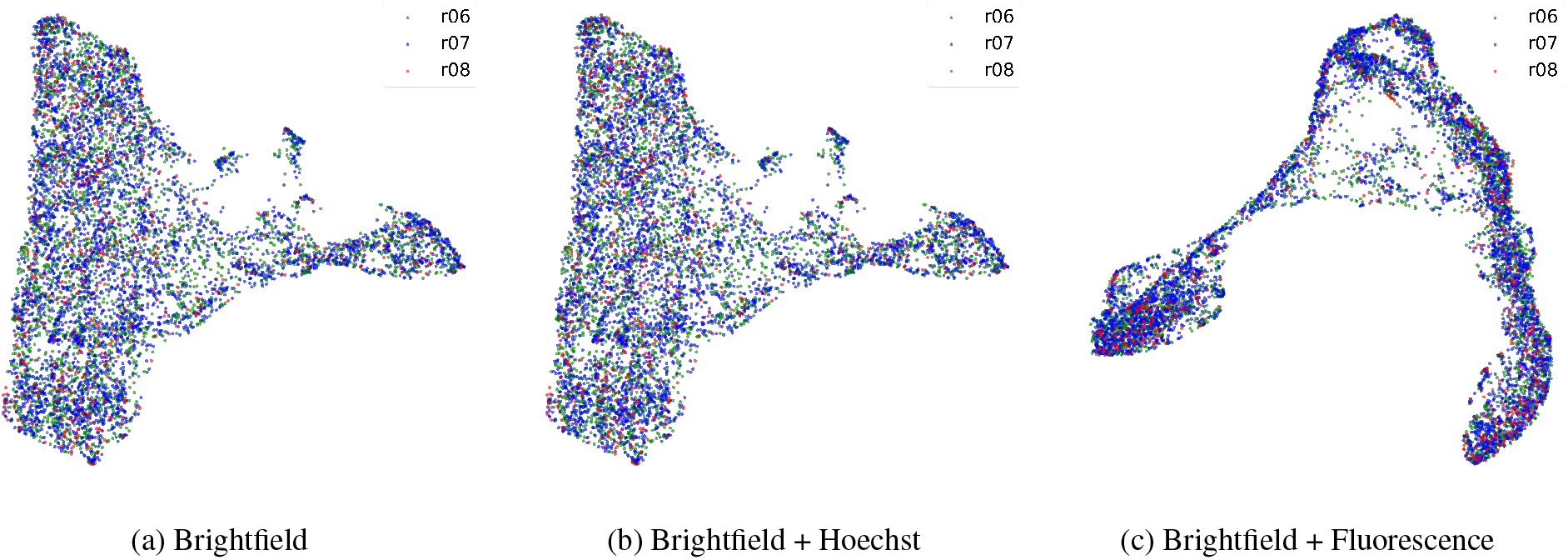
2D embeddings of cells from different patients in the dataset. The embeddings were calculated using UMAP from the feature map at the last layer before the last global average pooling in the network. (a) Embeddings from the model trained using only brightfield images. (b) Embeddings from the model trained with brightfield images and Hoeschst signal. (c) Embeddings from the model trained with brightfield images and all fluorescence signals.

**Figure 11:**
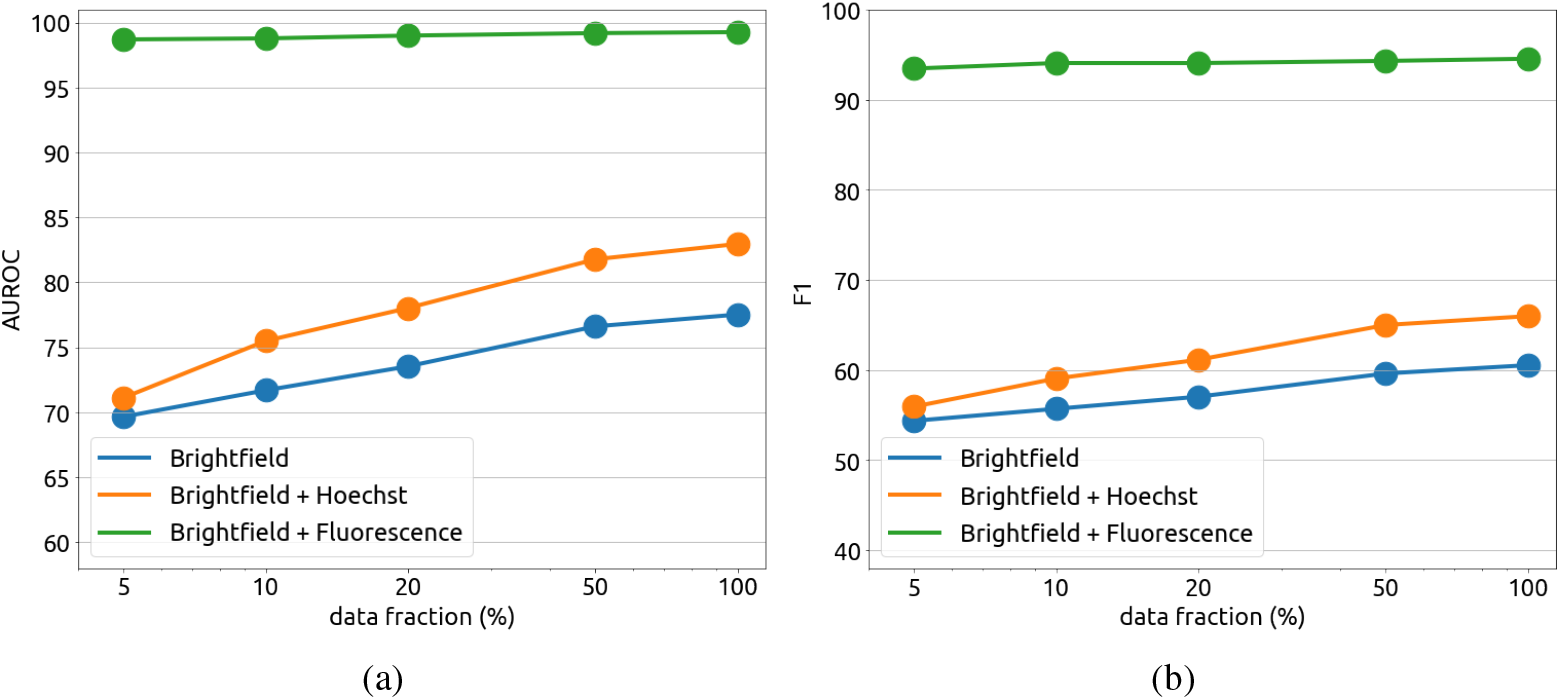
Impact of the training set size on the AUROC and F1 performances of cancer cell classification. Performances of the classifier were measured on the validation set. With full florescence signals as input, the model readily learned to identify cancer cells even with only a small data subset (green curve). In other settings, performances increased linearly as the data grew exponentially.

## Usage Notes

The detailed instruction for reproducing our work was described in the directory detection and classification of our Github.

## Code availability

All code used in this experiment was written in Python3 and could be publicly accessed at https://github.com/cmb-chula/CancerCellVision-CCA. The code is based on PyTorch(28) and MMDetection(22).

## Acknowledgements

This work was funded by the Asahi Glass Foundation (to S.S.), the Grant for Center of Excellence in Computational Molecular Biology, Ratchadapisek Sompoch Endowment Fund, Chulalongkorn University (to S.S. and E.C.), and the Second Century Fund (C2F), Chulalongkorn University (to C.K. and S.S.).

## Author contributions statement

N.I. conceived the experiments, P.I. conducted the experiments, C.P., C.K., and N.L. analyzed the results, Q.T., E.C., and S.S. supervised the research. C.P. and S.S. wrote the original draft. All authors reviewed and contributed to the revision of the manuscript.

## References

[1] Siddarth Rawal, Yu-Ping Yang, Richard Cote, and Ashutosh Agarwal. Identification and quantitation of circulating tumor cells. Annual Review of Analytical Chemistry, 10(1):321–343, 2017. doi: 10.1146/annurev-anchem-061516-045405. URL https://doi.org/10.1146/annurev-anchem-061516-045405. PMID: 28301753.

[2] Yue Ming, Yuan yuan Li, Haiyan Xing, Minghe Luo, Zi wei Li, Jianhong Chen, Jingxin Mo, and Sanjun Shi. Circulating tumor cells: From theory to nanotechnology-based detection. Frontiers in Pharmacology, 8, 2017.

[3] Petra Bankó, Sun Lee, Viola Nagygyörgy, Miklos Zrinyi, Chang Chae, Dong Cho, and András Telekes. Technologies for circulating tumor cell separation from whole blood. Journal of Hematology & Oncology, 12, 05 2019. doi: 10.1186/s13045-019-0735-4.

[4] Arun Satelli, Zachary Brownlee, Abhisek Mitra, Qing Meng, and Shulin Li. Circulating tumor cell enumeration with a combination of epithelial cell adhesion molecule- and cell-surface vimentin-based methods for monitoring breast cancer therapeutic response. Clinical chemistry, 61, 10 2014. doi: 10.1373/clinchem.2014.228122.

[5] Yan Xu, Biao Liu, Fengan Ding, Zhou Xiaodie, Pin Tu, Bo Yu, Yan He, and Peilin Huang. Circulating tumor cell detection: A direct comparison between negative and unbiased enrichment in lung cancer. Oncology Letters, 13, 04 2017. doi: 10.3892/ol.2017.6046.

[6] Zhongyi Zhu, Si Qiu, Kang Shao, and Yong Hou. Progress and challenges of sequencing and analyzing circulating tumor cells. Cell Biology and Toxicology, 34, 10 2018. doi: 10.1007/s10565-017-9418-5.

[7] Anca Ciurte, Cristina Selicean, Olga Soritău, and Rares Buiga. Automatic detection of circulating tumor cells in darkfield microscopic images of unstained blood using boosting techniques. PLoS ONE, 13, 2018.

[8] Carlos Aguilar-Avelar, Brenda Soto-García, Diana Aráiz, Juan Yee, Miguel Esparza, Franco Chacón, Jesus Delgado-Balderas, Mario Alvarez, Grissel Trujillo de Santiago, Lauro Gómez-Guerra, Liza Velarde-Calvillo, Alejandro Abarca, and J. Wong-Campos. High-throughput automated microscopy of circulating tumor cells. Scientific Reports, 9:1–9, 09 2019. doi: 10.1038/s41598-019-50241-w">10.1038/s41598-019-50241-w.

[9] Kai Yao, Nash Rochman, and Sean Sun. Cell type classification and unsupervised morphological phenotyping from low-resolution images using deep learning. Scientific Reports, 9:1–13, 09 2019. doi: 10.1038/s41598-019-50010-9.

[10] Claire Chen, Ata Mahjoubfar, Li-Chia Tai, Ian Blaby, Allen Huang, Kayvan Niazi, and Bahram Jalali. Deep learning in label-free cell classification. Scientific Reports, 6:21471, 03 2016. doi: 10.1038/srep21471.

[11] Eric M. Christiansen, Samuel J. Yang, D. Michael Ando, Ashkan Javaherian, Gaia Skibinski, Scott Lipnick, Elliot Mount, Alison O’Neil, Kevan Shah, Alicia K. Lee, Piyush Goyal, William Fedus, Ryan Poplin, Andre Esteva, Marc Berndl, Lee L. Rubin, Philip Nelson, and Steven Finkbeiner. In silico labeling: Predicting fluorescent labels in unlabeled images. Cell, 173(3):792–803.e19, 2018. ISSN 0092-8674. doi: https://doi.org/10.1016/j.cell.2018.03.040. URL https://www.sciencedirect.com/science/article/pii/S0092867418303647.

[12] Roger Brent and Laura Boucheron. Deep learning to predict microscope images. Nature Methods, 15, 10 2018. doi: 10.1038/s41592-018-0194-9.

[13] Sunyoung Park, Richard Ang, Simon Duffy, Jenny Bazov, Kim Chi, Peter Black, and Hongshen Ma. Morphological differences between circulating tumor cells from prostate cancer patients and cultured prostate cancer cells. PloS one, 9:e85264, 01 2014. doi: 10.1371/journal.pone.0085264.

[14] Shen Wang, Yuyuan Zhou, Xiaochen Qin, Suresh Nair, Xiaolei Huang, and Yaling Liu. Label-free detection of rare circulating tumor cells by image analysis and machine learning. Scientific Reports, 10, 07 2020. doi: 10.1038/s41598-020-69056-1.

[15] Leonie L. Zeune, Yoeri E. Boink, Guus van Dalum, Afroditi Nanou, Sanne de Wit, Kiki C. Andree, Joost F. Swennenhuis, Stephan A. van Gils, Leon W.M.M. Terstappen, and Christoph Brune. Deep learning of circulating tumour cells. Nature Machine Intelligence, 2(2):124–133, Feb 2020. ISSN 2522-5839. doi: 10.1038/s42256-020-0153-x. URL https://doi.org/10.1038/s42256-020-0153-x.

[16] Jarno Drost and Hans Clevers. Organoids in cancer research. Nature Reviews Cancer, 18, 04 2018. doi: 10.1038/s41568-018-0007-6.

[17] Francesco Amato, Colin Rae, Maria Giuseppina Prete, and Chiara Braconi. Cholangiocarcinoma disease modelling through patients derived organoids. Cells, 9(4), 2020. ISSN 2073-4409. doi: 10.3390/cells9040832. URL https://www.mdpi.com/2073-4409/9/4/832.

[18] Kentaro Wada LLC. Labelme: Image polygonal annotation with python, 2022. URL https://github.com/wkentaro/labelme.

[19] Shaoqing Ren, Kaiming He, Ross Girshick, and Jian Sun. Faster R-CNN: Towards real-time object detection with region proposal networks. In C. Cortes, N. Lawrence, D. Lee, M. Sugiyama, and R. Garnett, editors, Advances in Neural Information Processing Systems, volume 28. Curran Associates, Inc., 2015. URL https://proceedings.neurips.cc/paper/2015/file/14bfa6bb14875e45bba028a21ed38046-Paper.pdf.

[20] Kaiming He, Xiangyu Zhang, Shaoqing Ren, and Jian Sun. Deep residual learning for image recognition, 2015.

[21] J. Deng, W. Dong, R. Socher, L. Li, Kai Li, and Li Fei-Fei. Imagenet: A large-scale hierarchical image database. In 2009 IEEE Conference on Computer Vision and Pattern Recognition, pages 248–255, 2009. doi: 10.1109/CVPR.2009.5206848.

[22] Kai Chen, Jiaqi Wang, Jiangmiao Pang, Yuhang Cao, Yu Xiong, Xiaoxiao Li, Shuyang Sun, Wansen Feng, Ziwei Liu, Jiarui Xu, Zheng Zhang, Dazhi Cheng, Chenchen Zhu, Tianheng Cheng, Qijie Zhao, Buyu Li, Xin Lu, Rui Zhu, Yue Wu, Jifeng Dai, Jingdong Wang, Jianping Shi, Wanli Ouyang, Chen Change Loy, and Dahua Lin. Mmdetection: Open mmlab detection toolbox and benchmark, 2019.

[23] Zhuang Liu, Hanzi Mao, Chao-Yuan Wu, Christoph Feichtenhofer, Trevor Darrell, and Saining Xie. A convnet for the 2020s. In Proceedings of the IEEE/CVF Conference on Computer Vision and Pattern Recognition, pages 11976–11986, 2022.

[24] Mingxing Tan and Quoc V. Le. Efficientnet: Rethinking model scaling for convolutional neural networks, 2020.

[25] Gao Huang, Zhuang Liu, Laurens Van Der Maaten, and Kilian Q. Weinberger. Densely connected convolutional networks. In 2017 IEEE Conference on Computer Vision and Pattern Recognition (CVPR), pages 2261–2269, 2017. doi: 10.1109/CVPR.2017.243.

[26] Leland McInnes, John Healy, Nathaniel Saul, and Lukas Großberger. Umap: Uniform manifold approximation and projection. Journal of Open Source Software, 3(29):861, 2018. doi: 10.21105/joss.00861. URL https://doi.org/10.21105/joss.00861.

[27] Caroline Uhler and G.V. Shivashankar. Nuclear mechanopathology and cancer diagnosis. Trends in Cancer, 4: 320–331, 2018. doi: 10.1016/j.trecan.2018.02.009.

[28] Adam Paszke, Sam Gross, Francisco Massa, Adam Lerer, James Bradbury, Gregory Chanan, Trevor Killeen, Zeming Lin, Natalia Gimelshein, Luca Antiga, Alban Desmaison, Andreas Köpf, Edward Yang, Zach DeVito, Martin Raison, Alykhan Tejani, Sasank Chilamkurthy, Benoit Steiner, Lu Fang, Junjie Bai, and Soumith Chintala. PyTorch: An Imperative Style, High-Performance Deep Learning Library. Curran Associates Inc., Red Hook, NY, USA, 2019.

